# RPAP3 impairs TRBP-mediated translation of TAR-containing RNAs and FTSJ3 recruitment onto the HIV-1 genomic RNA

**DOI:** 10.64898/2026.11.16.760968

**Authors:** Camille Virciglio, Inès Lemberger, Léane Chevrolet, Xavier Manival, Mathieu Rederstorff

**Affiliations:** Université de Lorraine, CNRS, IMoPA, F-54000 Nancy, France

## Abstract

The interaction between TRBP and RPAP3, which is mutually exclusive from that between TRBP and Dicer, revealed a possible role for RPAP3 in the regulation of miRNA activity. Besides miRNA processing and/or function, TRBP is involved in promoting the expression of TAR-containing transcripts, and to impede innate immune system from sensing HIV-1 in infected cells by recruiting the cellular 2’-*O*-methyltransferase FTSJ3 onto the viral RNA. Here, we show that RPAP3 prevents the TRBP stimulatory effect on the translation of TAR-containing reporters, both *in vitro* and *in cellulo,* without affecting TRBP’s ability to bind to the TAR. We also observe that virion production is increased in RPAP3 depleted cells. Additionally, we show that TRBP directly recruits the methyltransferase FTSJ3, and that RPAP3 inhibits this ability. These observations open new avenues for possible novel therapeutics strategies aimed at hindering TRBP pro-viral functions, via novel regulation mechanisms involving RPAP3.

## INTRODUCTION

The trans<u>a</u>ctivating response RNA (TAR RNA) binding protein (TRBP or TARBP2) is a <u>d</u>ouble-<u>s</u>tranded RNA (dsRNA) binding protein first identified as a binding factor of the TAR RNA element of the <u>h</u>uman immunodeficiency <u>v</u>irus HIV-1 and 2 genomic RNAs (1–4). It is composed of two canonical A type ds<u>R</u>NA <u>b</u>inding <u>d</u>omains (dsRBDs) and a third B type dsRBD involved in the association to its protein partners, such as Dicer (5,6). TAR RNA elements are found in the long terminal repeats at both the 5’- and 3’-terminal extremities of HIV genomic RNA (R-U5 and U3-R). The 5’-end TAR RNA element secondary structure is a very stable hairpin structure and was shown to be involved in translation suppression of downstream regions preventing the proper ribosome recruitment at the cap (7,8). However, TRBP binding to TAR RNA relieves this inhibitory effect. It was even suggested that HIV-1 TAR dependent translation might be enhanced by TAR RNA processing through the TRBP-Dicer interaction (9). In addition, other proteins can bind the TAR RNA structure in order to improve translation. This is the case of the DEAD-box polypeptide 3 (DDX3), an ATP-dependent RNA helicase that can bind the HIV-1 5’ extremity and notably the TAR RNA element in order to promote HIV RNA translation (10–12). HIV RNA translation initiation is mainly cap-dependent but can use other alternative pathways, notably an internal ribosome entry site (IRES) located directly 3’ to the TAR element, which is used in cellular conditions where cap-dependent translation initiation is inhibited, in order to ensure virus replication (13). A second IRES is also located within the *gag* ORF (14,15).

Additionally, TRBP was shown to promote HIV infection by directly or indirectly inhibiting PKR activation, which is triggered by TAR RNA, and leads to global translation inhibition by phosphorylation of one subunit of the eukaryotic initiation factor 2 (2). More precisely, TRBP can inhibit PKR activity by direct interaction, which is even reinforced when TRBP is phosphorylated (3,4). PKR activity can also be impeded by TRBP indirectly, as TRBP can bind to PKR’s activator PACT (protein activator of the dsRNA dependent kinase PKR), the sequestration of which preventing its interaction with PKR (2).

Additionally, it was recently described that TRBP recruited the cellular 2’-*O*-methyl transferase FTSJ3 to HIV (16). FTSJ3 subsequently methylates the viral genome at several specific positions, which enables the virus to escape the host’s innate immune response.

TRBP is also involved in the miRNA biogenesis and function pathways, mainly as one of Dicer’s possible co-factors (5,17–19). TRBP enables Dicer substrates selection specificity and their accurate processing, which contributes to guaranteeing gene silencing specificity by generating the appropriate mature miRNAs.

On the other hand, RPAP3 is a member of the HSC70/HSP90 co-chaperone R2TP complex, where it regulates HSP activities and intervenes as a central component to enable other R2TP complex components associations (20,21). Within the PAQosome, the R2TP complex is involved in the assembly and maturation of several ribonucleoprotein particles, among which C/D and H/ACA boxes snoRNPs, or RNA polymerase II complex (20,22–27).

Previously, we have shown that the TPR1 domain of RPAP3 directly interacted with the 3^rd^ dsRBD of TRBP, in a Dicer mutually exclusive manner, while RPAP3 did not interact with the protein PACT (28). In RPAP3 depleted cells, we also observed increased RNA interference efficiency, suggesting a possible role for RPAP3 in regulating miRNA activity, possibly through its interaction with TRBP, via a yet unclear mechanism (29). Our structural study also enabled to identify and characterize a RPAP3 mutant that lost its ability to bind TRBP.

Here, we investigated whether RPAP3 might interfere with the TRBP related translation of TAR-containing RNAs and/or with the recruitment of FTSJ3 onto the viral RNA, and the possible impact on HIV replication. Using *in vitro* translation assays, we showed that increasing quantities of wild-type RPAP3 decreased TAR-dependent translation, while the RPAP3 D161 mutant we characterized unable to bind TRBP, did not. Next, we observed increased expression of a firefly luciferase reporter featuring a TAR element at the 5’-UTR in RPAP3 depleted cells compared to wild-type cells, while binding of TRBP onto the TAR-containing mRNA reporters was the same in both conditions. Additionally, we observed that p24 production was enhanced in the RPAP3 depleted cell line after transfection with a proviral clone, and that IRES-dependent translation was affected as well. Finally, we demonstrated that the interaction between TRBP and FTSJ3 was direct, and that it was obstructed by RPAP3. Altogether, RPAP3 appears as a promising novel actor possibly involved in the regulation of HIV replication, by modulating TRBP/TAR-dependent activities, such as translation or 2’-*O*-methylation of TAR-containing RNAs.

## MATERIALS AND METHODS

### Cell culture

HCT116 OsTIR1 and HCT116 OsTIR1 RPAP3-AID cells (28) were maintained in Mc’Coys medium in a humidified 5% CO_2_ incubator at 37°C. The medium is supplemented with 10% of fetal bovine serum, 10 U/ml of penicillin/streptomycin and 2.9 mg/ml of glutamine.

### Transfection

HCT116 OsTIR1 and HCT116 OsTIR1 RPAP3-AID were seeded at 3.10^5^ cells per well (6-well plates) for TAR-dependent translation, or at 2.10^6^ cells per 10 cm dishes for 2’-*O*-methylation analysis and RIP experiment. The next day, cells were induced with 500 µM of auxin and then transfected using <u>p</u>oly<u>e</u>thylenimine (PEI) with a 1:2 ratio (DNA:PEI): 3 µg of DNA for 6-well plates and 8 µg of DNA for 10 cm dishes.

### Plasmids and cloning

For protein production, TRBP, RPAP3-WT and RPAP3 D161A ORFs were cloned into homemade pnEA-3CH vector at the *5′-NdeI* and *3′-BamHI* sites (maps available upon request). The pGl2-TAR-Luc and the HIV molecular clone pNL4-3 Δenv plasmids were graciously obtained from A. Gatignol (McGill University, Montreal, CA) and J.C. Paillart (IBMC, Strasbourg, F), respectively.

### Antibodies

Antibodies and dilutions for Western Blot were the following: rabbit polyclonal anti-RPAP3 at 1:1000 dilution (Sigma-Aldrich, SAB1411438); polyclonal rabbit anti-TRBP at 1:1000 dilution (Abcam, ab72110); anti-HIV1 p24 antibody p24 (Abcam, ab32352) at 1:1000 dilution.

### Recombinant protein production and purification in *Escherichia coli*

*Escherichia coli* BL21 (DE3) pRARE2 cells were transformed with the pnEA-3CH vector and grown in LB medium. After reaching an OD of 0.7, the culture was induced with 100 µM of iso<u>p</u>ropyl-β-d-thio<u>g</u>alactopyranoside (IPTG) and then incubated overnight at 20°C. The culture was centrifuged at 4000 x g for 30 min at 4°C and bacteria were lysed with cold lysis buffer (25 mM HEPES, pH 7.5, 300 mM NaCl, 0.5 mM TCEP, 10 mM Imidazole) and sonicated (Output 2, Duty cycle: 50%, 30 sec on/ 30 sec off, 3 times). To remove any nucleic acids, a precipitation step with 0.0125% PEI was performed, followed by centrifugation at 4000 x g for 30 min at 4°C. The His-tagged protein was first purified by affinity chromatography using TALON resin for 30 min on a rotator followed by 3 washing steps. The protein was next eluted with the elution buffer (25 mM HEPES pH 7.5, 300 mM NaCl, 0.5 mM TCEP, 300 mM Imidazole) and selected elution fractions were injected onto an Aktä Purifier HPLC system (Superdex® 200 Increase 10/300 GL column), or a Q chromatography for TRBP, and eluted in a NaCl gradient buffer (10 mM HEPES pH 7.5, 300-1000 mM NaCl, 0.5 mM TCEP). Proteins were analyzed by SDS-PAGE, and their concentration quantified using a QuBit device prior to being aliquoted and stored at -20°C.

### *In vitro* transcription and translation

First, for *in vitro* transcription, the luciferase reporter pGL2 -TAR-Firefly was amplified by PCR using Phusion polymerase to get a linearized DNA template. Transcription was performed at 37°C for 2 h using the MEGAscript kit (Thermo Fisher Scientific) in a total volume of 20 µl containing 0.5 µg of linearized DNA template, 100 µM of dTNP and T7 RNA polymerase according to the manufacturer’s protocol. The transcribed RNA was then purified with the MEGAclear Transcription Clean-Up kit, dosed, aliquoted and stored at -80°C before experiments. RNA reporter was translated in rabbit reticulocyte lysates (Promega L4960). Briefly, one translation reaction was made in a final volume of 25 µl containing 1 µg of RNA, 17.5 µl of Rabbit Reticulocytes, 70 µM of leucine and methionine and 0.5 U of RNase inhibitor, together with the recombinant protein at 30°C for 90 min. Translation is then quantified by measuring the luciferase luminescence with a luminometer by adding luciferase substrate (Promega).

### Translation *in cellulo*

Cells were transfected using PEI with 3 µg of reporter’s plasmid and/or plasmid expressing TRBP or RPAP3 WT ORFs. Cells were harvested the next day in the <u>P</u>assive <u>L</u>ysis <u>B</u>uffer (PLB) buffer from the Promega Luciferase Assay kit, lysed by 3 freeze/thaw cycles and were centrifuged. 10 µl of the supernatant were used in a 96-plates to measure luminescence using 100 µl of Firefly substrate. Data were normalized using the mRNA reporter level quantified by RT-qPCR.

### RT-qPCR

HCT116 OsTIR1 and HCT116 OsTIR1 RPAP3-AID cells were cultured in 10 cm Petri dishes (seeded at 2 × 10^6^ cells per dish) and transfected with 8 µg of pGL2 TAR-Firefly reporter. Total RNAs were extracted 24 h post-transfection with TRIzol reagent (Thermo Fisher Scientific), and resuspended in RNase-free water. 2 µg of RNA were treated with DNase RQ1 for 1 h at 37°C and then the reverse transcription was performed with MMLV enzyme using random hexamers and polydT primers for 1 h. cDNAs at a dilution of 1/4 were then used to perform qPCR with Supermix iTaq Universal SYBR Green reagent using appropriate primer (Firefly forward: ACTTCGAAATGTCCGTTCGG; Firefly reverse: GCAACTCCG-ATAAATAACGCG; Gag forward: AAAAATTTTGACTAGCGGAGGC ; Gag reverse: AGGCCAGGATTAACTGCGAATC). The plate was loaded into a thermal cycler according to the following PCR program: hot start : 10 min at 95°C, 40 amplification cycles: 15 s at 95°C, 15 s at 60°C, 15 s at 72°C, terminal elongation step : 10 min at 70°C; melt curve step 30 s at 95°C, 1 min at 60°C then go to 95°C with a 0.5°C/s slope.

### RNA immunoprecipitation

Cells were harvested 24 h after auxin induction and pGL2-TAR Firefly reporter transfection. Cells were lysed for 10 min at 4°C in HTNG Buffer (20 mM HEPES, pH7.9, 150 mM NaCl, 1 mM MgCl2, 1 mM EDTA, 1% Triton X-100, 10% glycerol) and centrifuged to pellet debris. RIP experiment was performed using 30 µl of Dynabeads Protein G (Invitrogen), coated with 4 µg of antibody against TRBP for 1 h and incubated with HCT116 OsTIR1 and HCT116 OsTIR1 RPAP3-AID lysates on a rotator for 2 h. Beads were then washed 3 times with HTNG 1% Buffer and then RNA was eluted in a final volume of 100 µl of HTNG 1% containing 1% SDS and 5 µl of a PK protease (20 mg/ml), in a thermoshaker at 1200 rpm for 30 minutes at 37°C. RNA were extracted using TRIzol LS, phenol/chloroform/isoamyl alcohol, precipitated, ethanol-washed and resuspended in RNase-free water. Total RNA was treated with RQ1 DNase, reverse-transcribed using MMLV enzyme and cDNA were used to perform a qPCR using appropriate primers (Forward: ACTTCGAAATGTCCGTTCGG ; Reverse : GCAACTCCGATAA-ATAACGCG).

### Virus production

The day after HCT116 OsTIR1 and HCT116 OsTIR1 RPAP3-AID cells were seeded in plates, cells were induced with 500 µM of auxin and transfected with 8 µg of HIV molecular clone (pNL4-3 Δenv) using the PEI transfection protocol. After 48 h, cells were harvested and virions production from cell extracts was monitored by western-blotting or quantified by HIV p24 titration by ELISA (Abcam).

### TRBP-FTSJ3 Pull Down

His-tagged FTSJ3 and CRD-TRBP proteins were produced as previously described (30,31). His-tagged FTSJ3 produced in a cell-free system was then first dialyzed for 2 h at 4°C in pull-down buffer (25 mM HEPES, 200 mM NaCl, 10 mM imidazole, 10% glycerol, 1× protease inhibitor cocktail). All subsequent steps were performed at 4°C. Three pull-down conditions were tested: His-FTSJ3 alone, CRD-TRBP alone, and His-FTSJ3 with CRD-TRBP. For each condition, proteins were incubated with 100 µL of TALON resin slurry and either 500 µL of cell-free extract or 20 µg of purified CRD-TRBP for 30 min with gentle agitation. The resin was then centrifuged at 700 × g for 3 min and washed three times with 200 µL of pull-down buffer. For the His-FTSJ3/CRD-TRBP interaction assay, 20 µg of CRD-TRBP was added to resin-bound His-FTSJ3 and incubated for an additional 30 min. The resin was washed three times with 200 µL of pull-down buffer and bound proteins were eluted with 25 µL of Laemmli buffer. Samples were then loaded on a 10% SDS-PAGE gel and subjected to both Coomassie blue staining and Western Blot analysis to assess the interaction between the two proteins. Flag-TRBP IPs were performed as previously described (28).

## RESULTS

### RPAP3 impairs TAR dependent translation *in vitro*

TRBP stimulates HIV replication by inhibiting PKR activation, by promoting escape from the host’s immune system or by improving translation of viral mRNAs (7,16,32). To investigate the possible regulatory role RPAP3 might play during TRBP dependent translation, we measured expression of a firefly luciferase reporter construct featuring the TAR element at its 5’-end, using *in vitro* translation in rabbit reticulocyte lysates (7) **(Fig. 1, Supp. Fig. 1 and 2)**. In our experiments, as expected, growing concentrations of TRBP (0.33-1.33 µM) improved translation efficiency, up to 2-fold with the 1.33 µM condition **(Supp. Fig. 1A)**. Higher concentrations inhibited translation, as previously observed (7). We hypothesized that RPAP3, by interacting with TRBP, could interfere with its stimulatory effect on translation. Indeed, increasing concentrations of RPAP3 (3.33-10 µM) reduced translation of the firefly reporter both in presence of 1.33 µM of TRBP **(Fig. 1A)** or in reticulocyte lysates only **(Supp. Fig. 1B)**. The effect observed is specific, as the RPAP3 D161A mutant, that does not bind TRBP (28), did not alter translation efficiency of the same reporter **(Fig. 1B, Supp. Fig. 1C)**. This suggests that RPAP3 interferes with the translation of the reporter, possibly by impeding TRBP’s translation stimulatory effect, but even appears to reinforce the inhibitory effect on translation caused by the TAR RNA structure.

**Figure 1.**
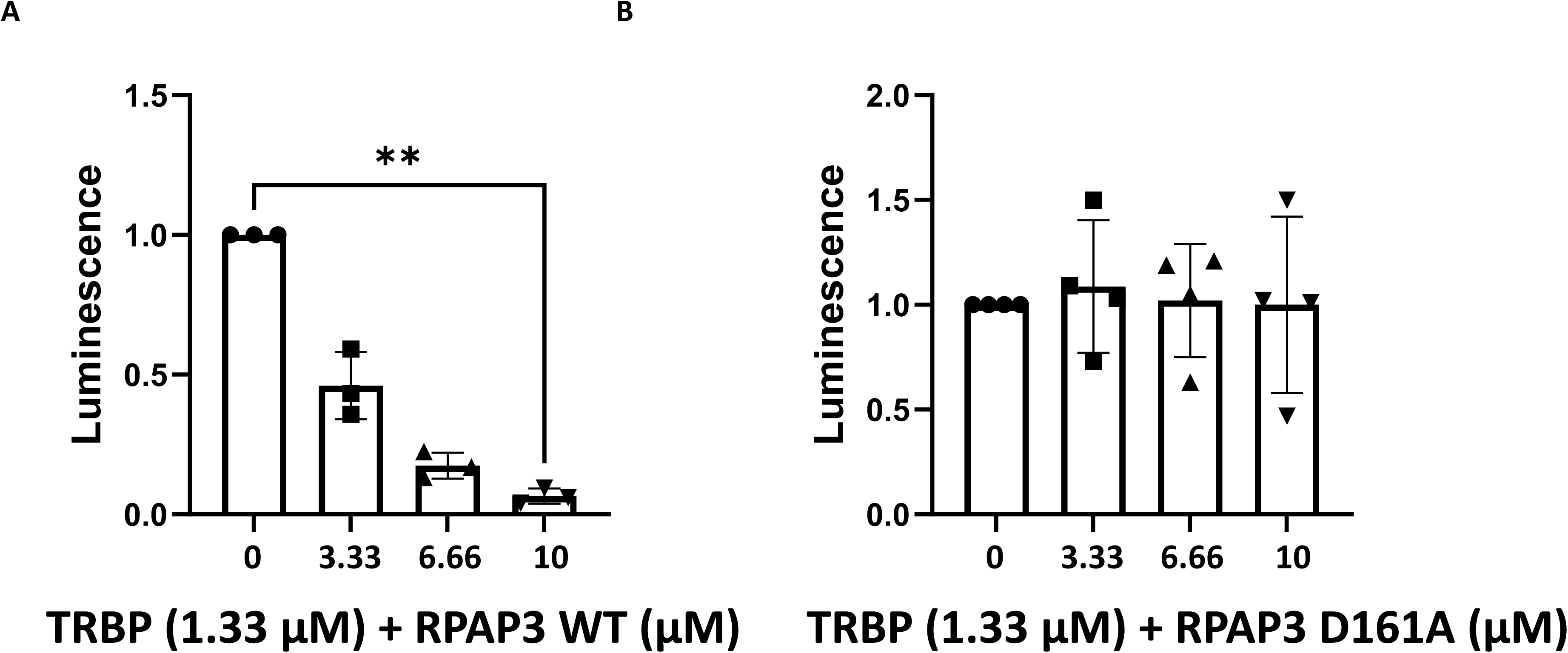
RPAP3 impairs the expression of TAR-containing luciferase transcripts *in vitro*. TAR-Firefly reporter mRNAs were translated in rabbit reticulocyte lysates in the presence of a constant concentration of His-TRBP (1.33 µM) and by adding increasing concentrations of RPAP3 WT (A) or RPAP3 D161A (B) (0, 3.33, 6.66 and 10 µM). Data were analyzed using the Kruskal-Wallis statistical test and were normalized to the HCT116 OsTIR1 cells. Individual points and means ± SD are shown. n = 3 (A) and n= 4 (B) biological replicates.

**Figure 2.**
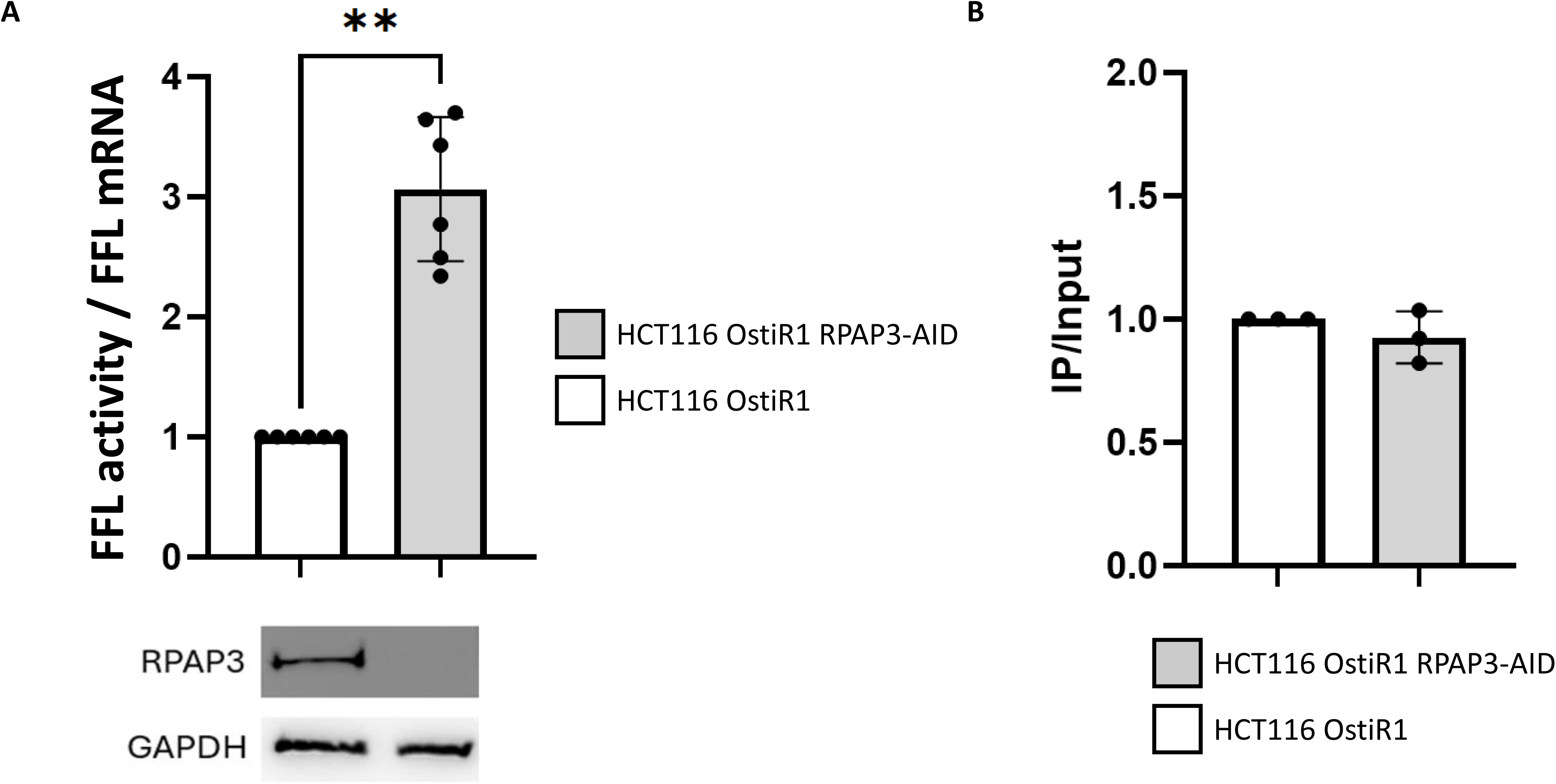
Depletion of RPAP3 leads to an increase in TAR-dependent translation. (A) TAR-dependent translation quantification in HCT116 OsTIR1 control cells and HCT116 OsTIR1 RPAP3-AID (RPAP3 depleted) cells. Cells were transfected with the pGL2 CMV-T7 TAR-luciferase plasmid. After 24 h of incubation, cells were washed twice in phosphate-buffered saline. One-tenth of the cells were used for luciferase analysis, and the remainder was processed for RNA extraction and qRT-PCR analysis on both the transfected luciferase and control GAPDH mRNAs. Individual points and means ± SD are shown. Data were normalized to the HCT116 OsTIR1 cells and analyzed using the Mann-Whitney statistical test. n = 6 biological replicates. (B) Quantification of the association of TRBP with TAR-containing transcripts. HCT116 OsTIR1 and HCT116 OsTIR1 RPAP3-AID were transfected with 8 µg of the pGL2 CMV-T7 TAR-luciferase plasmid. Binding of TRBP to the TAR element was quantified by RNA immunoprecipitation experiments followed by qRT-PCR using specific primers. n=3 biological replicates. Individual points and means ± SD are shown. Data were normalized to the HCT116 OsTIR1 cells and analyzed using the Mann-Whitney statistical test.

### TAR-dependent translation is improved in RPAP3 depleted cells

We next investigated whether this inhibitory effect could be observed *in cellulo,* using our HCT116 cell model system in which RPAP3 can be depleted owing to a conditional inducible degron strategy (28). To do so, we transiently transfected in both control and RPAP3-depleted cell lines a firefly luciferase reporter featuring the TAR element at its 5’-end **(Supp. Fig. 2A)**. As previously observed, translation efficiency of the firefly luciferase reporter was much less efficient with the TAR-containing RNA reporter constructs, in both cell lines **(Supp. Fig. 2B)**. In these cells, translation of the TAR-containing construct was improved by 3 folds in RPAP3 depleted cells (HCT116 OsTIR1 RPAP3-AID) compared to controls cells (HCT116 OsTIR1) **(Fig. 2A)**, which is in concordance with the observations we made using *in vitro* translation, where increasing quantities of RPAP3 led to the reduction of the translation efficiency.

### TRBP efficiently binds the TAR RNA in presence of RPAP3

As the binding of RPAP3 could interfere with TRBP’s association to the TAR element, we measured by RNA pull-down assays the amounts of TAR containing RNAs that co-precipitated with TRBP. Binding of TRBP to the TAR RNAs was unaltered in both conditions, suggesting that the interaction of RPAP3 with TRBP does not impact its association onto the RNA **(Fig. 2B)**. This is in adequation with our hypothesis that RPAP3 may, once associated to TRBP onto the TAR element, prevent its stimulatory effect on the translation of the reporter, rather than impeding its association to the RNA.

### p24 production is increased in RPAP3 depleted cells

Translation of HIV mRNAs contributes to viral proteins production, virions assembly, secretion and subsequent infection of new cells. The TAR element is present at the 5’ end of the HIV genomic RNA, and of all spliced viral mRNAs, where it is associated to an Internal Ribosome Entry Site (IRES) **(Supp. Fig. 3A)**. Even though IRES dependent initiation of translation is minor, compared to cap dependent translation (14,33), we wanted to measure the RPAP3 impact on translation efficiency in presence of the full 5’ region (TAR + IRES), in a more physiological context. We therefore wanted to assess whether RPAP3 had an impact on virions synthesis in our cell lines. To do so, p24 synthesis and virions production were quantified by measuring p24 levels in cells transfected with the proviral molecular clone pNL4-3 Δenv, both in cell extracts (intra cytoplasmic p24) and in cell culture supernatants (virions) **(Fig. 3, Supp. Fig. 3B)**. We observed a visible increase in both p55 and p24 expression levels in the HCT166 OsTIR1 RPAP3 AID cells, as assessed by western-blotting **(Fig. 3A)**. This observation was confirmed by ELISA quantification of p24, which is 3 times more abundant in RPAP3 depleted cells compared to the control cells **(Fig. 3B)**. The same observation is made when quantifying p24 expression from virions isolated and purified from cell culture medium, suggesting an increased virion production in the absence of RPAP3 **(Supp. Fig. 3B)**. Therefore, RPAP3 negatively regulates virion production, likely via TRBP and TAR-dependent translation mechanisms, among other potential effects.

**Figure 3.**
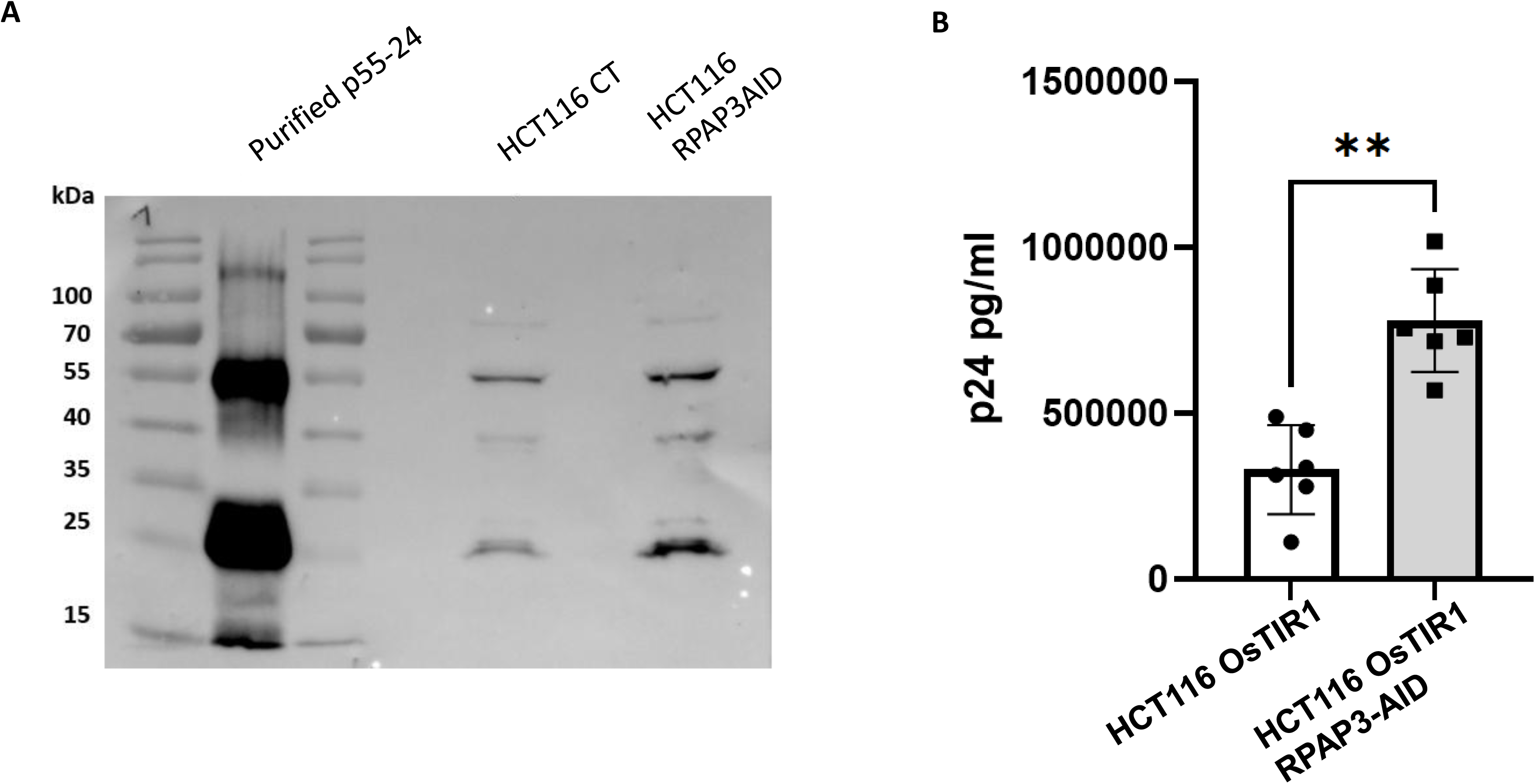
Virus production is increased in the absence of RPAP3. HCT116 OsTIR1 and HCT116 OsTIR1 RPAP3-AID cells were transfected with the HIV molecular clone pNL4-3 Δenv. Cells were harvested and lysed 48 h post-transfection and cytoplasmatic p24 production was analyzed by Western blot (A) or ELISA quantification (B). Western Blot shows a representative result from one biological replicate. Individual points and means ± SD are shown. Data were analyzed using the Mann-Whitney statistical test. n = 6 biological replicates.

We also wanted to test the impact of RPAP3 when translation was strictly IRES dependent, using a bicistronic gene reporter system **(Supp. Fig 3A, right)**. As expected, no firefly luciferase was expressed in the absence of the IRES element at the 5’ of the firefly reporter gene, while presence of the IRES enabled translation **(Supp. Fig 3C).** Again, we observed that firefly luciferase expression was improved in RPAP3 depleted cells compared to wild-type cells **(Supp. Fig 3D)**, suggesting that IRES dependent translation was affected by RPAP3 as well.

### TRBP directly interacts with FTSJ3

It was previously shown that TRBP recruited the methyltransferase FTJS3 to 2’-*O*-methylate the HIV genomic RNA at least at 17 positions (16), however, the nature of the association between the two proteins, direct or indirect, was not determined. To answer this question, we first expressed and purified both recombinant TRBP and FTSF3 protines *in vitro*. Next, we performed a pull-down experiment, using the purified His-Tagged version of full-length FTSJ3, and a purified full-length TRBP featuring a CRD-Tag. His-FTSJ3 efficiently co-precipitated TRBP, proving that the interaction between FTSJ3 and TRBP is direct **(Fig. 4A)**.

**Figure 4.**
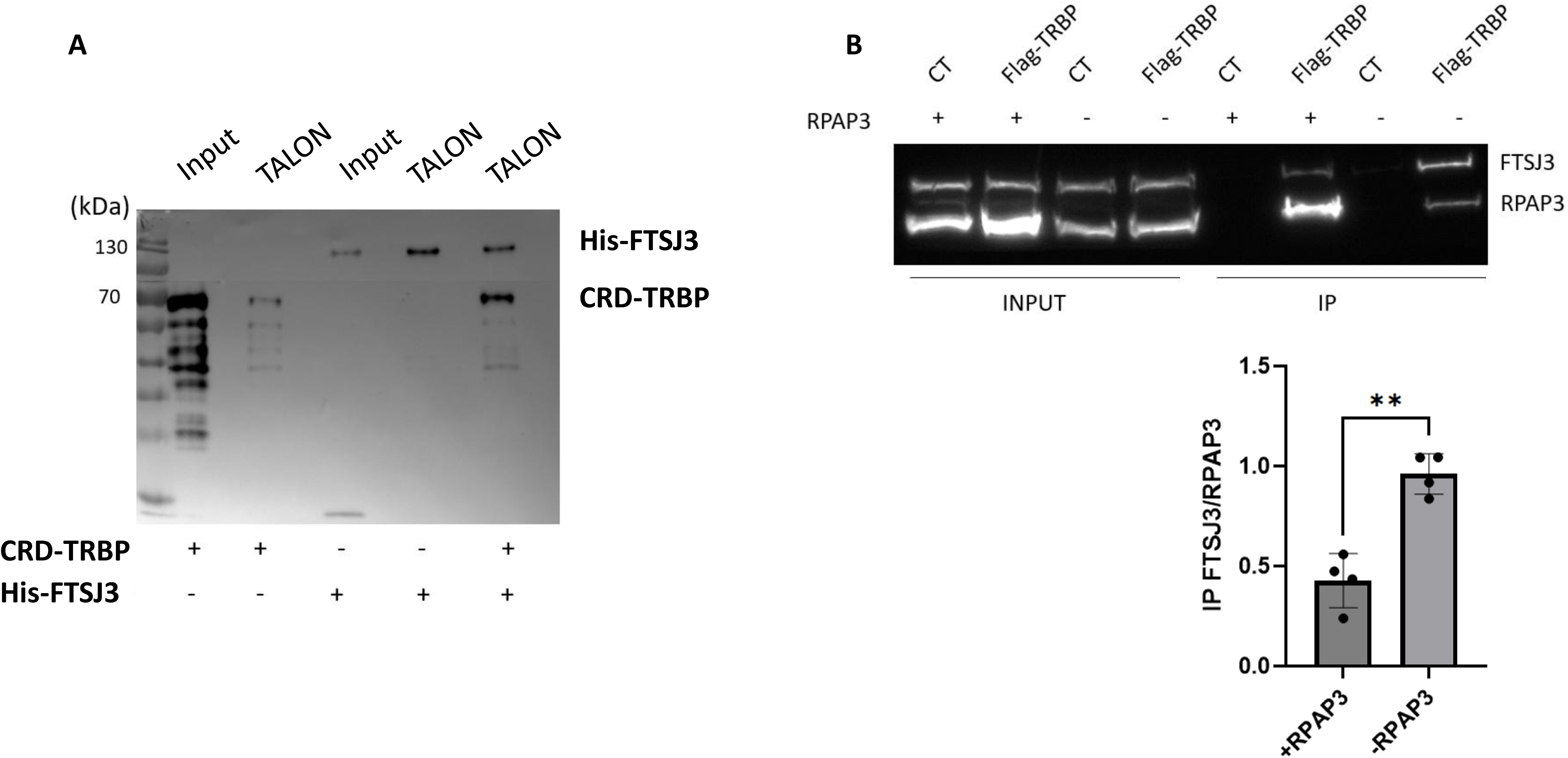
RPAP3 impairs the association between TRBP and FTSJ3. (A) Pull-down experiment performed employing *in vitro* produced and purified recombinant TRBP and FTSJ3 proteins. HIS-tagged FTSJ3 was bound to TALON beads and incubated in presence of purified CRD-TRBP, revealing a direct interaction between FTSJ3 and TRBP. (B) Co-immunoprecipitation of FTSJ3 and RPAP3 using a stable cell line expressing an inducible Flag-TRBP protein, transiently overexpressing RPAP3 or not. Relative ratios of FTSJ3/RPAP3 precipitated (IP) by Flag-TRBP are represented as a histogram below the Western-Blot. Data were analyzed using a Welch t-test. Individual points and means ± SD are shown. n = 4.

### RPAP3 impairs the association between TRBP and FTSJ3

TRBP interacts with its protein partners via its non-canonical dsRBD3. Like the mutually exclusive interactions we identified between TRBP and RPAP3 on one hand, and between TRBP and Dicer on the other hand (28), involving the dsRBD3 of TRBP, we wanted to investigate whether RPAP3 could also interfere with the recruitment of FTSJ3 by TRBP. To do so, in a cell line expressing a Flag-tagged version of TRBP, we immunoprecipitated both FTSJ3 and RPAP3, in conditions where RPAP3 was transiently overexpressed or not **(Fig. 4B)**. As observed by western-blotting, TRBP efficiently co-precipitated both RPAP3 and FTSJ3 **(Fig. 4B, upper panel)**. However, the ratios of RPAP3 and FTSJ3 precipitated by TRBP are inverted when RPAP3 is overexpressed **(Fig. 4B, lanes 6 and 8)**, with about two times more FTSJ3 precipitated in the absence of RPAP3 **(Fig. 4B, lower panel)**, suggesting again a competitive mutually exclusive association of TRBP either with RPAP3 or FTSJ3.

## DISCUSSION

TRBP mostly improves HIV-1 replication by inhibiting PKR-dependent antiviral response, but also via direct interaction with the TAR element, affecting translation or post-transcriptional modification of the viral RNAs (2,7,16). Here, we explored the impact of the R2TP component RPAP3, which binds TRBP in a mutually exclusive manner from Dicer, on the translation stimulation of TAR-containing RNAs, as well as on the recruitment of the methyl-transferase FTSJ3.

As observed in the course of miRNA function, where we showed an increase in miRNA activity in the absence of RPAP3 (28), here we observed the increased translation of a reporter protein downstream of the TAR regulatory element, while increasing amounts of RPAP3 lead to the opposite effect. This was observed not only during *in vitro* transcription/translation assays, but also *in cellulo*. Interestingly, our RPAP3 D161A mutant, which does not enable interaction with TRBP, did not reproduce the effect *in vitro*. We also observe that these effects are not linked to a decreased association of TRBP to the TAR element, as, both in presence or absence of RPAP3, TRBP binding to the TAR-containing reporter mRNA was identical. This was expected, as it is the dsRBD3 of TRBP that is involved in binding its protein partners, including RPAP3 (28,34), while the dsRBD1 and 2 are involved in RNA binding (Fig. 5).

**Figure 5.**
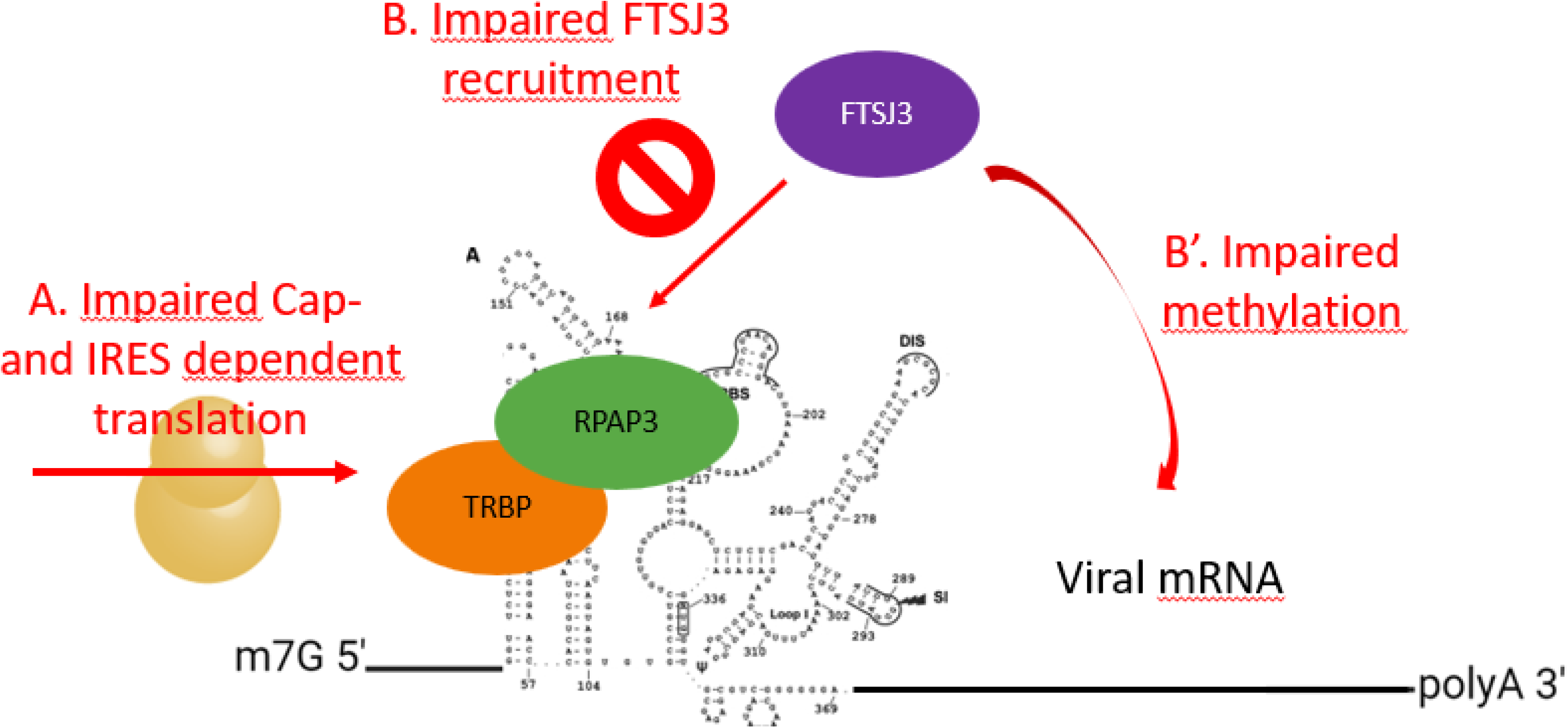
Model of RPAP3-mediated inhibition of TRBP-dependent translation and FTSJ3 recruitment on HIV-1 genomic RNA. TRBP binds the TAR-element via its dsRBD1 and 2, while RPAP3-bound TRBP loses its stimulatory effect on translation, leading to reduced cap-/IRES-dependent translation. Additionally, RPAP3 competes with FTSJ3 for binding to TRBP, via TRBP’s dsRBD3. This competition prevents FTSJ3 recruitment, leading to impaired 2’-*O*-methylation. Altogether, both RPAP3-dependent mechanisms ultimately decrease viral protein synthesis and virion production.

In a more physiological system, we similarly observed that in cells transfected with the proviral clone pNL4-3 Δenv, antigen protein p24 and virion production was increased about 3-fold in RPAP3 depleted cells, indicating that the RPAP3 negative impact on translation not only affected TAR-containing reporters, but viral mRNA featuring the entire 5’-LTR as well, including the IRES element located just downstream of the TAR. Even though translation of viral mRNA is mostly cap-dependent, presence of the IRES therefore does not seem to reduce RPAP3’s negative impact on translation. We even showed that RPAP3 had a similar impact on translation when initiation was strictly IRES dependent, using a bicistronic reporter. This is surprising as it shows that RPAP3 impairs both cap-dependent and independent translation, via TRBP and the TAR. Indeed, while RPAP3 impairs the stimulatory effect of TRBP on cap-dependent translation, the IRES dependent translation, with the IRES located downstream of the TAR, should not be affected. Our explanation for that is that presence of RPAP3, once recruited by TRBP in the TAR vicinity, might lead to a strong sterical constrain, preventing ribosome access or progression even on the IRES (Fig. 5). However, the mechanism by which RPAP3 precisely blocks translation of downstream regions is not yet deciphered. We previously showed that TRBP immunoprecipitated not only RPAP3, but the whole R2TP complex, with the interaction between RPAP3 and TRBP being direct (28). Presence of a large complex such as the R2TP would of course even more contribute to impair ribosome progression.

How does TRBP exactly destabilize the TAR region to increase translation is not clear. It is possible that TRBP contributes to recruiting helicases, such as DDX3 for instance, to unwind the TAR and downstream sequences to facilitate translation. RPAP3, once bound to TRBP, might interferes with these mechanisms, by avoiding recruitment of the proteins involved. *Komori et al.* even suggested that the TAR was destabilized by being cleaved by Dicer (9). As the interaction between TRBP and Dicer, and between TRBP and RPAP3, are mutually exclusive, the fact that RPAP3 might prevent TRBP from recruiting Dicer and therefore avoid cleavage of the TAR could be another hypothesis to explore.

As RPAP3 is involved in various functions, even though the effects we observe seem specific, one cannot exclude that other, TRBP unrelated, pathways, might be involved, including transcription regulation via RNA polymerase 2 assembly, which relies on RPAP3 (26). TRBP is also known to modestly affect non-TAR reporters (7).

Finally, modification of the viral RNA constitutes a key feature in viral replication (16,35). TRBP was shown to recruit the cellular 2’-*O*-methyltransferase FTSJ3 onto the genomic RNA, via the TAR element (16). We showed here that the interaction between the two partners was direct, and even though we lack structural data at this stage, it is likely that it involves the dsRBD3 domain of TRBP (Fig. 5). Moreover, we showed that RPAP3 limited the recruitment of FTSJ3 by TRBP, and thus, might decrease virus escape from the immune system, by reducing the 2’-*O*-methylation of the viral genome, which we are currently investigating.

Altogether, RPAP3 therefore constitutes a promising new player in regulation of HIV replication. Studying the RPAP3/TRBP interaction might be a promising unprecedented research track for novel therapeutics strategies at different levels, while TRBP has already proven interesting therapeutic potential in the past (36).

## ACKNOWLEDGMENTS

We wish to thank Anne Gatignol and her lab, as well as Jean-Christophe Paillart, for the gracious gift of several plasmids. We thank the Team 1 of the IMoPA unit, as well as Jean-Christophe Paillart and Franck Martin for helpful discussions. We thank Marie-Eve Chagot and Lionel Imbert for their help regarding protein production. We thank Virginie Marchand and Yuri Motorin, as well as the EpiRNA-seq platform. This work was supported by the ANRS grants [20413 AO 2020–2 CSS 11, ECTZ133498] and [AO 2022-1 CSS 11, ECTZ188028].

---

**Supplementary Figure 1. RPAP3 impairs the expression of TAR-containing luciferase transcripts *in vitro*.** TAR-Firefly reporters mRNAs were translated in rabbit reticulocyte lysates in presence of increasing concentrations of TRBP (0, 0.33, 0.66 and 1.33 µM) (A), RPAP3 WT (B) or RPAP3 D161A (C) (0, 3.33, 6.66 and 10 µM). Data were analyzed using the Kruskal-Wallis statistical test and were normalized to the HCT116 OsTIR1 cells. Individual points and means ± SD are shown. n = 5 (A, C) and 6 (B) biological replicates.

**Supplementary Figure 2. TAR-dependent translation.** (A) Luciferase reporter featuring the TAR element at its 5’-end and controls. (B) Consistent with established literature (7), the presence of the TAR element strongly decreased translation efficiency of the firefly luciferase reporter in both HCT116 OsTIR1 (black) and HCT116 OsTIR1 RPAP3-AID (grey).

**Supplementary Figure 3. IRES-dependent translation.** (A) Right: pNL4-3 derived GAG mRNA featuring the full 5’-LTR region, including the TAR element and the IRES; Left: dual luciferase reporter featuring the full 5’-LTR upstream the firefly luciferase. Translation of Renilla is under dependence of the 5’ cap, and presence of the EMCV IRES ensures ribosome drop-off after translation of the Renilla luciferase. (B) HCT116 OsTIR1 and HCT116 OsTIR1 RPAP3-AID cells were transfected with the HIV molecular clone pNL4-3 Δenv. Cell culture was stopped 48 h post-transfection and virions were prepared from cell culture media for p24 quantification by ELISA. Individual points and means ± SD are shown. Data were analyzed using the Mann-Whitney statistical test. n = 6 biological replicates. (C) Presence of the IRES is mandatory for the expression of the firefly reporter. (D) Depletion of RPAP3 in the HCT116 OsTIR1 RPAP3-AID cell line leads to an increase in IRES-dependent translation.

## Notes

### Competing Interest Statement

The authors have declared no competing interest.

