## supplementary figures 1-3 for "RPAP3 impairs TRBP-mediated translation of TAR-containing RNAs and FTSJ3 recruitment onto the HIV-1 genomic RNA"

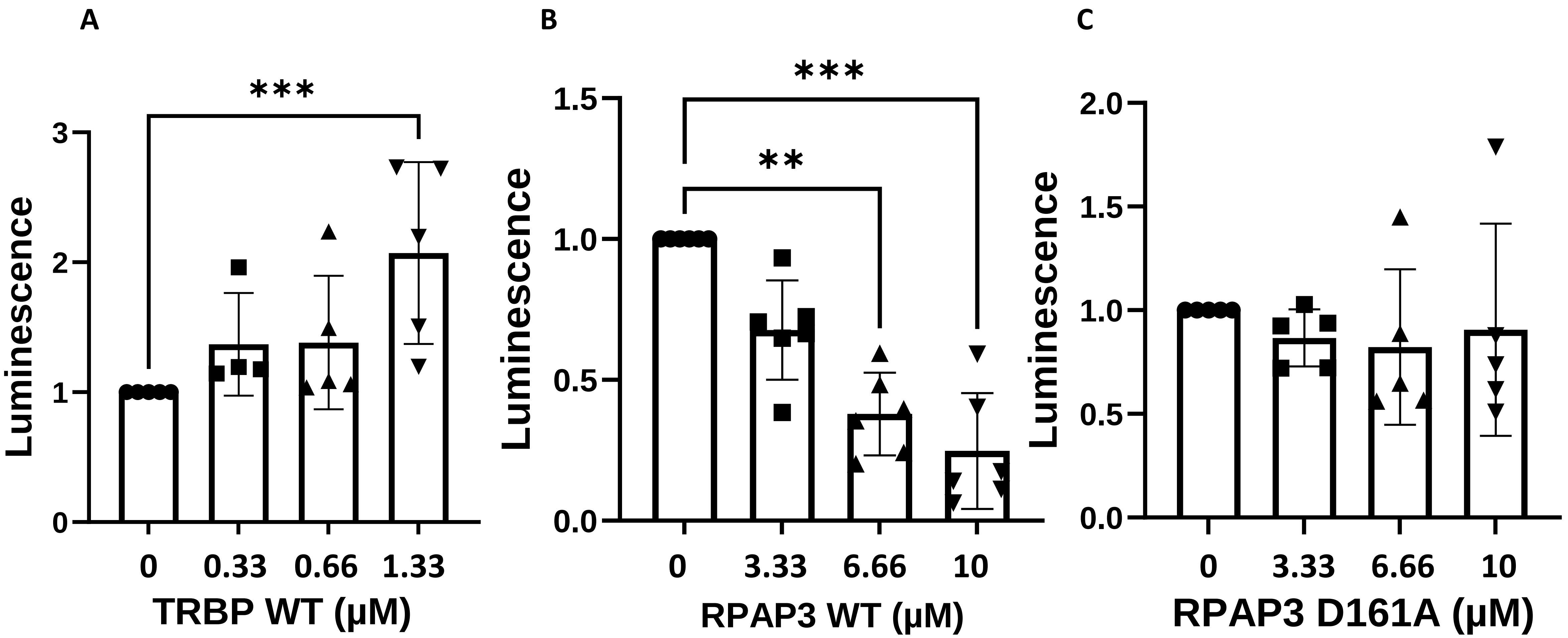
A
B
C
***
***
**
Luminescence
Luminescence
Luminescence
3
2
1
0
1.5
1.0
0.5
0.0
2.0
1.5
1.0
0.5
0.0
0
0.33
0.66
1.33
TRBP WT (μM)
0
3.33
6.66
10
RPAP3 WT (μM)
0
3.33
6.66
10
RPAP3 D161A (μM)

Supp. Fig. 1

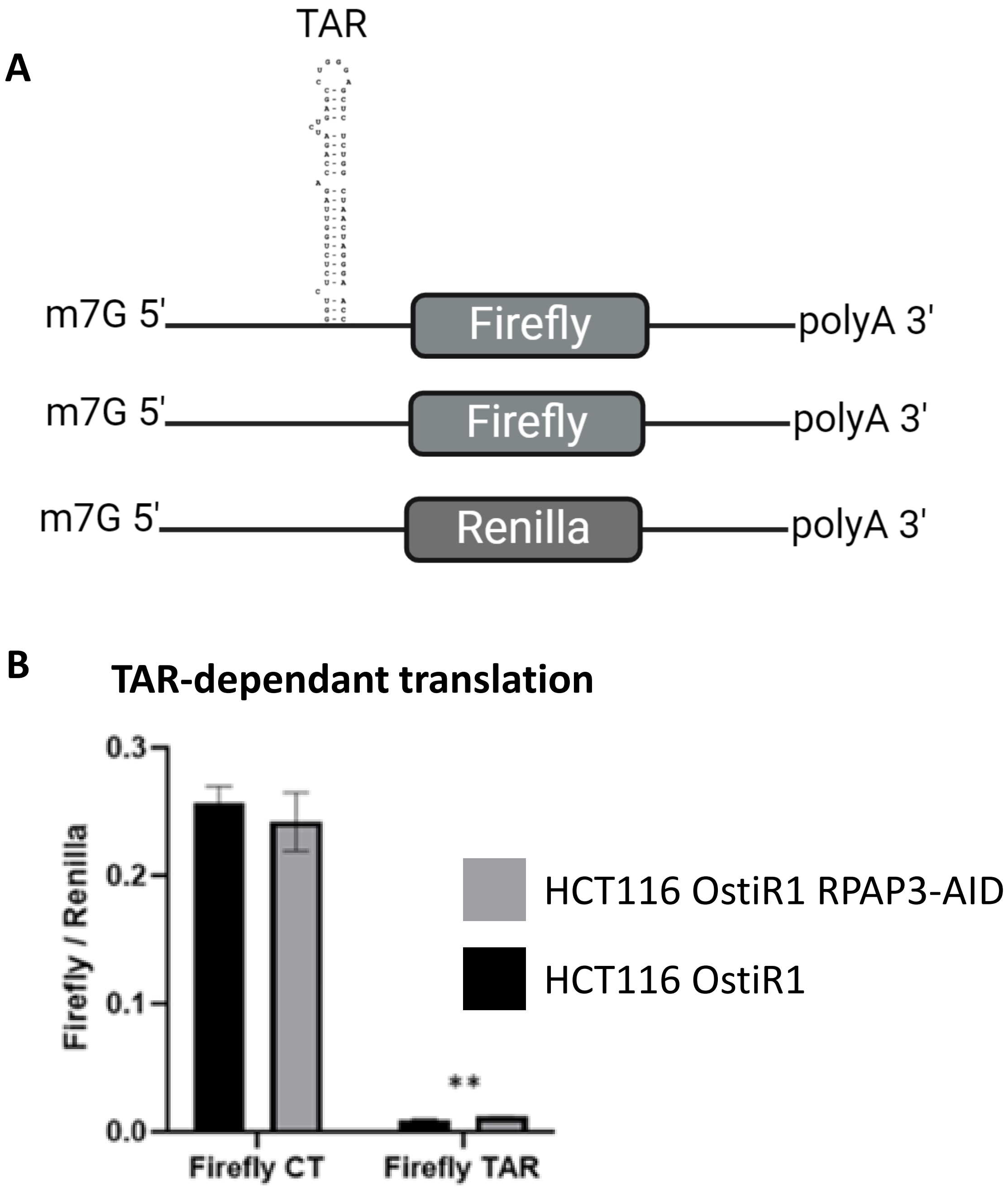
A
TAR
m7G 5'
Firefly
polyA 3'
m7G 5'
Firefly
polyA 3'
m7G 5'
Renilla
polyA 3'
B
TAR-dependant translation
Firefly / Renilla
0.3
0.2
0.1
0.0
**
Firefly CT
Firefly TAR
HCT116 OstiR1 RPAP3-AID
HCT116 OstiR1

**Supp. Fig. 2**

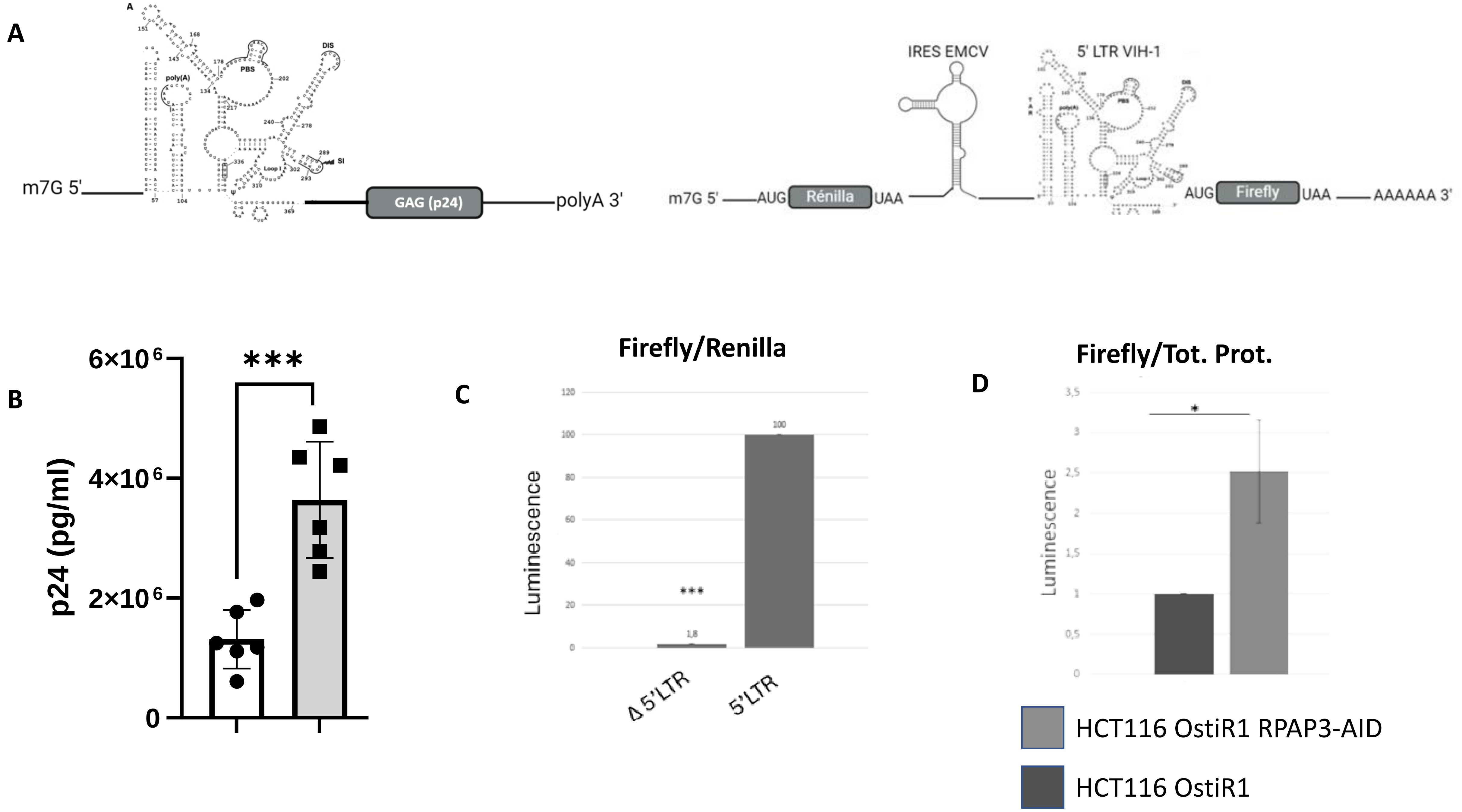
A
m7G 5'
GAG (p24)
polyA 3'
IRES EMCV
5' LTR VIH-1
m7G 5'
AUG
Rénilla
UAA
AUG
Firefly
UAA
AAAAAA 3'
B
p24 (pg/ml)
6×10^6
4×10^6
2×10^6
0
***
C
Firefly/Renilla
Luminescence
***
1,8
100
Δ 5'LTR
5'LTR
D
Firefly/Tot. Prot.
Luminescence
*
HCT116 OstiR1 RPAP3-AID
HCT116 OstiR1

Supp. Fig. 3
